# Melanopsin contributes to circadian photic responses in mice in a sex-dependent manner

**DOI:** 10.64898/2025.12.10.693461

**Authors:** Kayla C. Miguel, Marcos L. Aranda, Jacob D. Bhoi, Matthew Diaz, Jennifer A. Evans, Tiffany M. Schmidt

**Affiliations:** Department of Neurobiology, Northwestern University, Evanston, IL, USA; Department of Ophthalmology, Feinberg, School of Medicine, Northwestern University, Chicago, IL, USA; Department of Biomedical Sciences, Marquette University, Milwaukee, WI, USA

## Abstract

Proper entrainment of the body’s circadian rhythms to the environment is critical to human health. Light is one of the strongest cues driving circadian photoentrainment of the central circadian pacemaker, the suprachiasmatic nucleus (SCN), via projections from the melanopsin-expressing intrinsically photosensitive retinal ganglion cells (ipRGCs). Circadian research has historically centered males, and recent work has revealed multiple sex-differences in circadian circuitry and function, indicating that our understanding of this system in females is severely limited. Moreover, while recent studies have investigated the role of hormonal modulation of light responses, the additional possibility that ipRGC inputs may also be sex-dependent has not been directly tested. Here, we report that not only do ipRGCs in female mice show higher levels of melanopsin expression, but that melanopsin also plays a larger role in shaping circadian photic responses in females compared to males. Collectively, these results define a new retinal source for sex-dependent differences in circadian behavior.

## Introduction

Circadian rhythms are regulated by a number of internal and environmental cues, with light being one of the most important. Most mammals synchronize their behavior to the day/night cycle and employ behavioral strategies to re-synchronize their activity to changes in the timing of light. These circadian behavioral responses to light are important because circadian disruptions, including those resulting from mistimed light, can result in various negative health outcomes.^1–6^ However, much of our historical understanding of the circadian system has come from studies of males due to concerns about female hormone-driven variability.^7^ Consequently, despite decades of research on how light modulates circadian behaviors, more recent studies that include females have uncovered unexpected sex differences,^8,9^ revealing a major gap in our understanding of circadian circuits.

Female mice have been shown to have larger light-induced phase delays to a light pulse in the early evening and faster re-entrainment to a 6-hour simulated jet lag advance.^10–14^ These behavioral differences have been largely attributed to circulating estrogens, as they are decreased or abolished in gonadectomized mice as well as in estrogen receptor (ESR1) knockout mice. Indeed, estrogen has been shown to increase the excitability of neurons in the central circadian pacemaker, the suprachiasmatic nucleus (SCN),^15^ which could result in larger SCN responses to light signals. However, it is also possible that the light signals the SCN receives are themselves sex-dependent, but this possibility has yet to be directly tested.

Light input to the SCN arises solely from the melanopsin-expressing, intrinsically photosensitive retinal ganglion cells (ipRGCs).^16,17^ ipRGCs integrate synaptic rod and cone input with their own intrinsic melanopsin photoresponse ^17–19^ and are necessary for photic circadian responses.^20–22^ Despite the new evidence that photic circadian responses are sex-dependent, the role of ipRCGs in these differences has not been examined. Here, we set out to determine whether ipRGCs exhibit sex-dependent differences, and whether these differences might also contribute to reported behavioral differences. We find that, compared to males, female mice have higher expression of melanopsin in ipRGCs and show more severe deficits in light-driven circadian behaviors following deletion of the melanopsin gene.

## Results and discussion

### Female ipRGCs have higher *Opn4* expression

Given the central role of melanopsin (encoded by the *Opn4* gene) in light-driven circadian behaviors and ipRGC function, we first sought to compare *Opn4* expression levels in male versus female mice. To do this, we used RNAscope fluorescent in situ hybridization to label *Opn4* mRNA in single ipRGCs of whole mount male and female Opn4^Cre/+^ retinas. Interestingly, female ipRGCs showed significantly higher levels of daytime *Opn4* expression (Figure 1A, p = 0.0207, Mann-Whitney).

**Figure 1.**
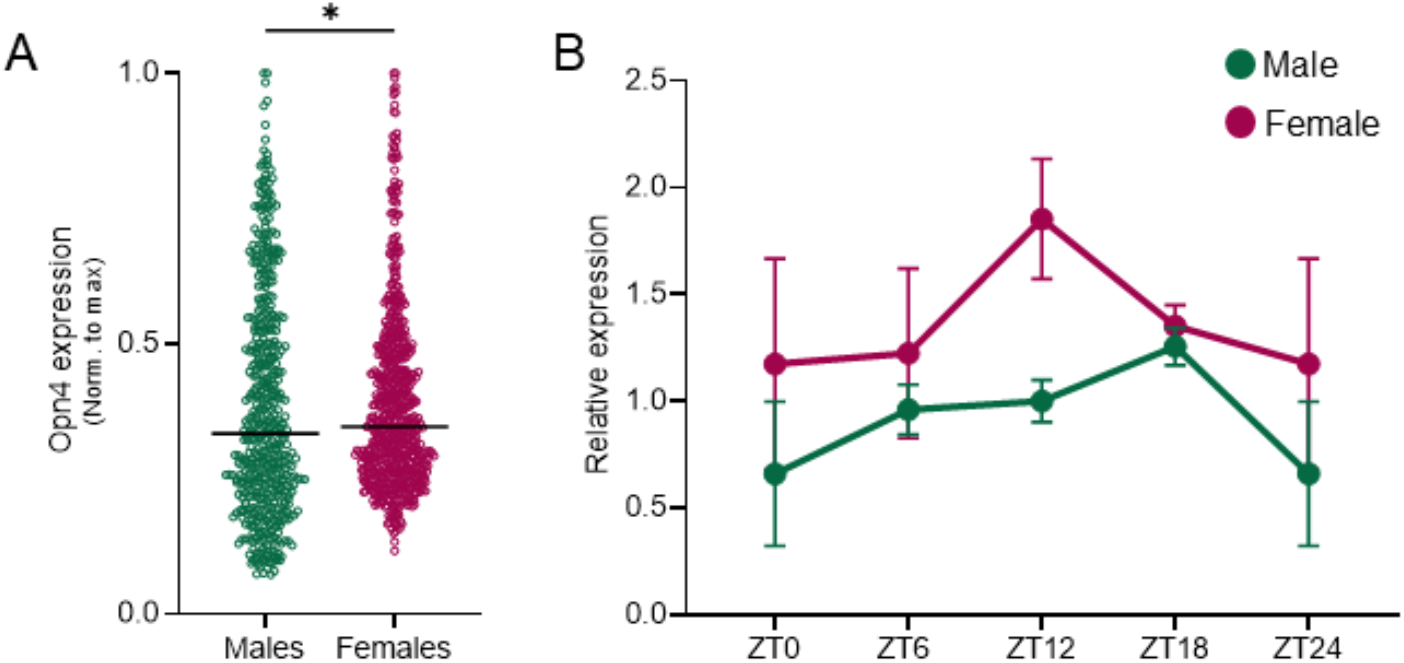
Female ipRGCs have higher *Opn4* expression. (A) Average pixel intensity per ipRGC of *Opn4* mRNA using RNAscope in male and female whole retinas (male: n = 512; female: n = 690; *p = 0.0207, Mann-Whitney). (B) Relative *Opn4* mRNA expression measured by qPCR normalized to female ZT 0 average. Error bars indicate ± SEM, n = 3-4 mice per group per time point. Data for ZT0 are double plotted at ZT24.

Because *Opn4* expression is rhythmic over the course of the day,^23,24^ we next compared *Opn4* mRNA levels in males and females at different times of day. To do this, we performed qPCR to quantify *Opn4* mRNA expression in whole mouse retinas from male and female Opn4^Cre/+^ mice across 4 different time points (ZT0, ZT6, ZT12, ZT18). *Opn4* mRNA expression was rhythmic in both male and female mice, peaking at ZT12 for females and ZT18 for males (Figure 1B). In agreement with our RNAscope analysis, we found that females had consistently higher *Opn4* mRNA expression across the day, with the most marked difference at ZT12. Collectively, these findings indicate that ipRGCs in the female retina have higher *Opn4* mRNA expression than ipRGCs in the male retina, suggesting that these differences may contribute to previously reported sex differences in light-driven circadian behavior.

### Sex-dependent impacts of melanopsin on the early evening photic phase delay

Compared to males, female mice show larger photic phase delays to a light pulse given in the early evening (CT14-16) and express higher levels of *Opn4* (Figure 1A).^10,11^ Intriguingly, this *Opn4* expression difference peaks at ZT12, just prior to the time window for the peak behavioral difference in photic phase delays (Figure 1B), suggesting that *Opn4* expression differences could play a role. To test this, we measured the phase shift of female and male Opn4^Cre/+^ (control) and melanopsin null, Opn4^Cre/Cre^ (MKO) littermates to a 15-minute, 1000-lux light pulse delivered at CT14 on the second day after mice were released into constant darkness (Figure 2A). If increased melanopsin drives the stronger phase delay in females, then the sex difference in phase delay magnitude should be abolished in melanopsin null mice. Though at this light intensity we only observed a trend toward a greater female photic phase delay in controls, we noted that this trend disappeared in in MKO male and female littermates, which showed phase delays similar to male controls (Figure 2B, Two-way ANOVA; Sex, F(1, 29) = 3.324, p = 0.0786; Genotype, F(1, 29) = 1.862, p = 0.1829).

**Figure 2.**
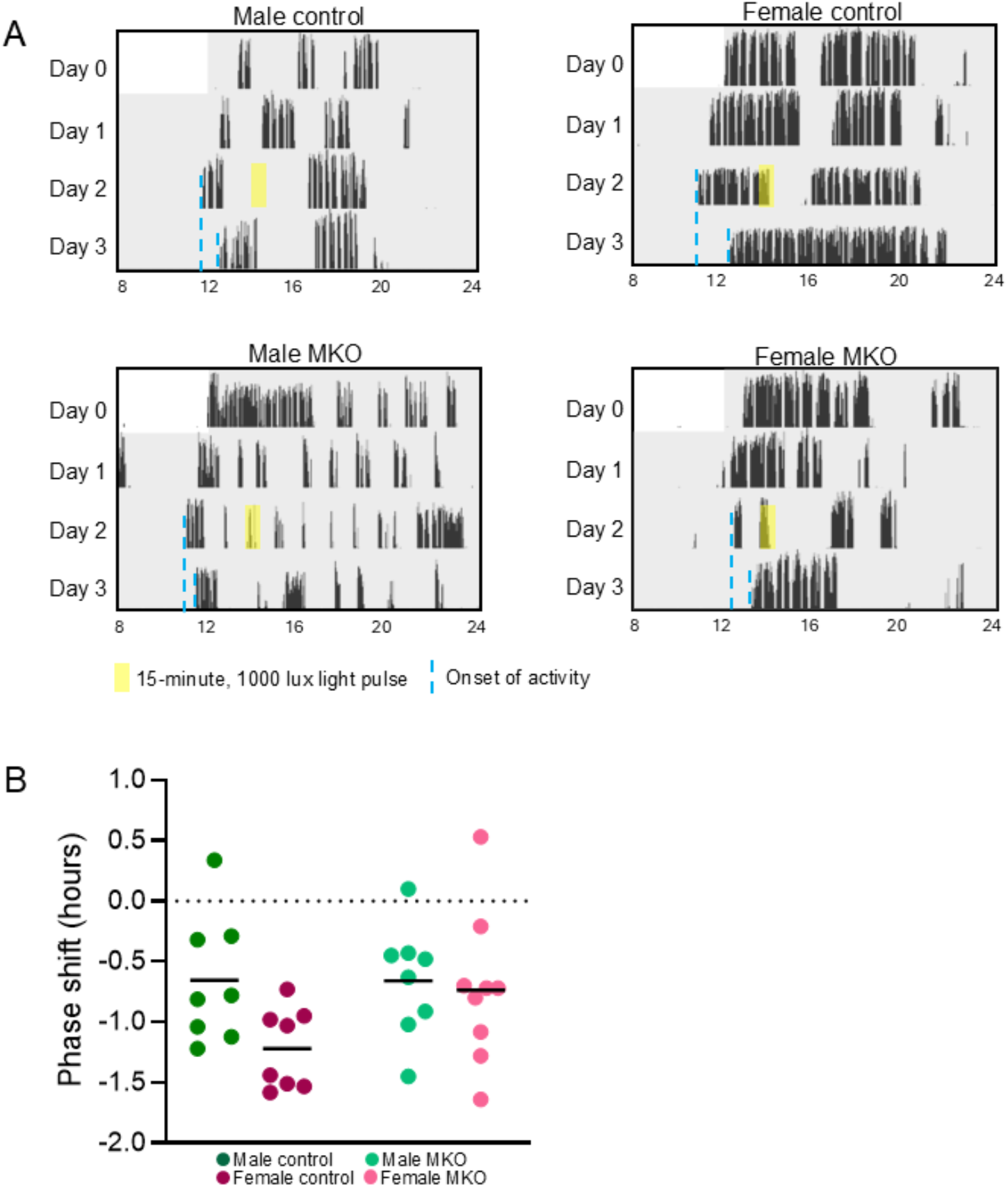
Sex-dependent impacts of melanopsin on the early evening photic phase delay at 1000 lux. (A) Example actograms with light pulse (yellow) and measured pre- and post-pulse onsets (dotted blue lines) represented. (B) Phase shift in hours after a 15-minute, 1000-lux light pulse at CT14 on the second day of constant dark (Two-way ANOVA; Sex, F(1, 29) = 3.324, p = 0.0786; Genotype, F(1, 29) = 1.862, p = 0.1829).

We reasoned that the lack of significant differences observed in male and female control mice could be due to the melanopsin-saturating, 1000 lux intensity of the light pulse. Importantly, this was a higher intensity than used in other studies that reported this sex difference. Therefore, we performed a similar experiment using a lower intensity light stimulus of 150-lux. We measured the phase shift of female and male control and MKO littermates to a 1-hour, 150-lux light pulse delivered at CT16 on the third day after mice were released into constant darkness. At this intensity, female control mice showed a significantly larger phase shift than male control littermates, similar to the trends observed at 1000 lux (Figure 3A-B, Two-way ANOVA; Sex, F(1, 39) = 8.998, p = 0.0047; Genotype, F(1, 39) = 11.55, p = 0.0016. Šídák’s multiple comparisons test; Control Female vs Control Male, p = 0.0048; Control Female vs MKO Male, p = 0.0008; Control Female vs MKO Female, p = 0.0020). This sex difference was absent in MKO mice, with female MKO mice showing a significantly reduced phase shift compared to female controls that was similar to male control and MKO littermates. These findings indicate that melanopsin is required for the larger early evening photic phase delay in females and also demonstrate that loss of melanopsin does not impact the magnitude of this phase delay in males.

**Figure 3.**
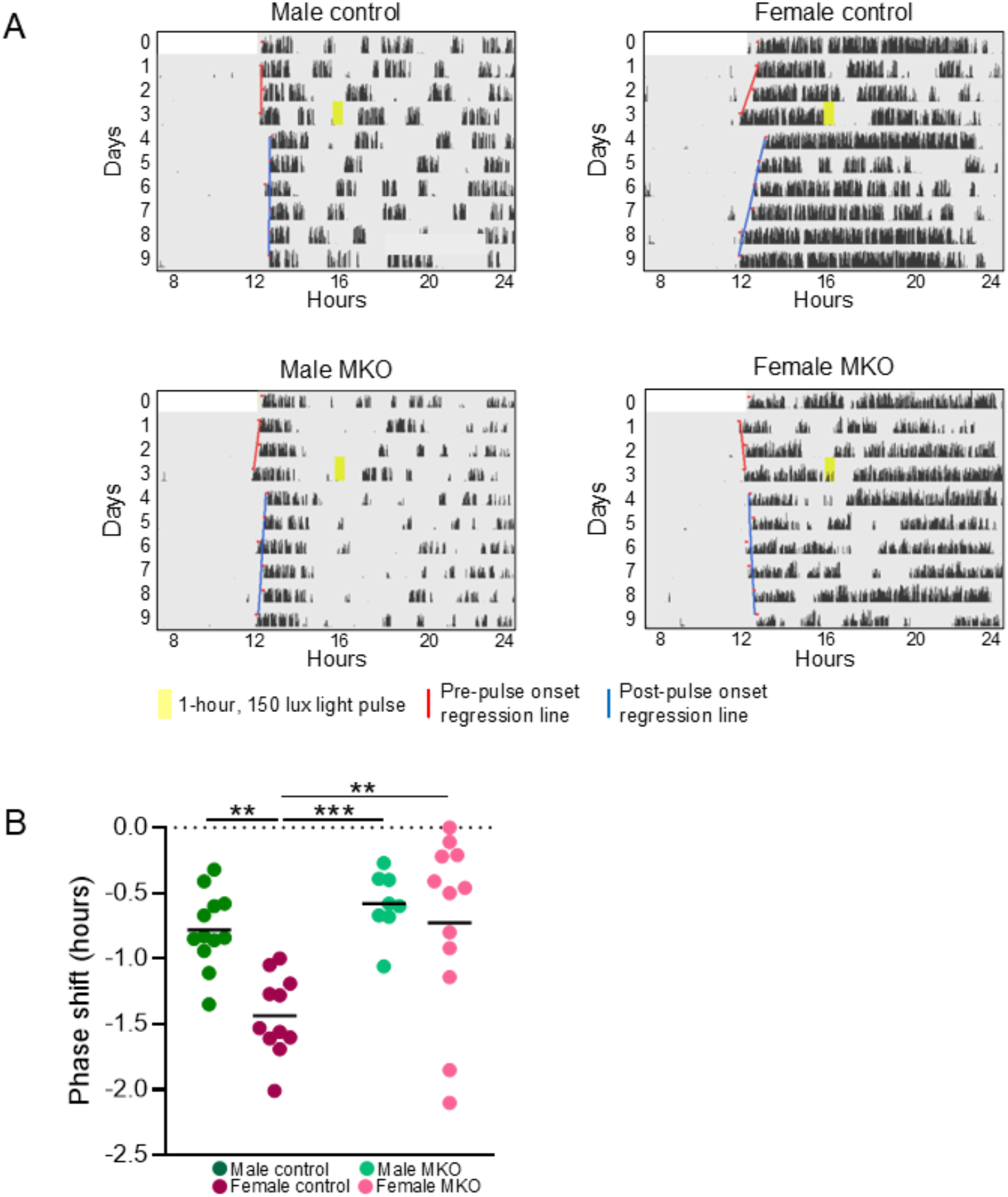
Sex-dependent impacts of melanopsin on the early evening photic phase delay at 150 lux. (A) Example actograms with light pulse (yellow), as well as pre-pulse (red) and post-pulse (blue) activity onset regression lines shown. (B) Phase shift in hours after a 1-hour, 150-lux light pulse at CT16 on the third day of constant dark. (Two-way ANOVA; Sex, F(1, 39) = 8.998, p = 0.0047; Genotype, F(1, 39) = 11.55, p = 0.0016. Šídák’s multiple comparisons test; Control Female vs Control Male, p = 0.0048; Control Female vs MKO Male, p = 0.0008; Control Female vs MKO Female, p = 0.0020).

Importantly, MKO and control littermates of both sexes showed similar, normal photoentrainment to a 100 lux 12:12 LD light schedule and re-entrainment to a 6-hour phase advance (Supplemental Figure 1A). While previous studies have shown that female mice are able to re-entrain faster than male mice to jet lag advances, ^12,14^ these studies used higher intensity light (∼250-350 lux). It is therefore possible that differences might be detectable at higher light levels. There is also evidence to suggest that rate of re-entrainment is estrus-stage dependent, with females re-entraining faster when they are in proestrus. ^13^ However, we did not track estrus stage in this study to avoid disrupting wheel running behavior with daily vaginal lavage.

### Lack of melanopsin results in a sex-specific deficit in negative masking

Mice decrease locomotor activity when presented with a light pulse in the early evening, and this behavior is referred to as negative masking. Mice lacking melanopsin show deficits in negative masking, indicating that melanopsin is important for this behavior.^25,26^ However, whether male and female mice have differences in negative masking has not been directly tested. Given that negative masking light pulses are typically given in early evening when the male/female difference in *Opn4* mRNA expression is largest (Figure 2B), it is possible that melanopsin may differentially impact negative masking in males versus females. We therefore sought to examine first whether male and female negative masking differs and next whether it is differentially impacted by melanopsin. To test this, we compared the sensitivity of the negative masking response of male and female control and MKO littermates to 3-hour light pulses of 100 or 300 lux, delivered between ZT14-17.

At 100 lux, both male and female control mice showed robust negative masking for the entire 3-hour light pulse, evidenced by a decrease in wheel revolutions during this period (Figure 4A-C). However, MKO male and female mice failed to sustain negative masking for the duration of the light pulse, with female MKO animals showing a more severe deficit compared to male MKO littermates (Figure 4B, Two-way ANOVA; Sex, F(1, 23) = 1.309, p = 0.2644; Genotype, F(1, 23) = 29.67, p < 0.0001, Šídák’s multiple comparisons test; Control Female vs MKO Male, p = 0.0486; Control Female vs MKO Female, p = 0.0004; Control Male vs MKO Female, p = 0.0003). At 300 lux, control male and female mice and MKO males showed similar negative masking for the entire 3 hour light pulse, while female MKO animals showed impaired negative masking, with only a transient suppression of wheel running activity in the first hour (Figure 4D-F) (Figure 4E, Two-way ANOVA; Sex, F(1, 19) = 6.652, p = 0.0184; Genotype, F(1, 19) = 7.206, p = 0.0147, Šídák’s multiple comparisons test; Control Male vs MKO Female, p = 0.0028; Control Female vs MKO Female, p = 0.0108; MKO Male vs MKO Female, p = 0.0072). The continued impairment of negative masking in female MKO animals at a higher light intensity indicates that melanopsin enhances negative masking in females at this intensity, while rod/cone photoreception in males compensates for the loss of melanopsin. Thus, despite similar degrees of negative masking in male and female controls, the photoreceptive systems and underlying retinal mechanisms driving these responses differs by sex. These results highlight that similar circadian behavioral output does not necessarily indicate that the underlying mechanisms are identical and suggest that additional sex differences in circadian circuits and behaviors are yet to be found.

**Figure 4.**
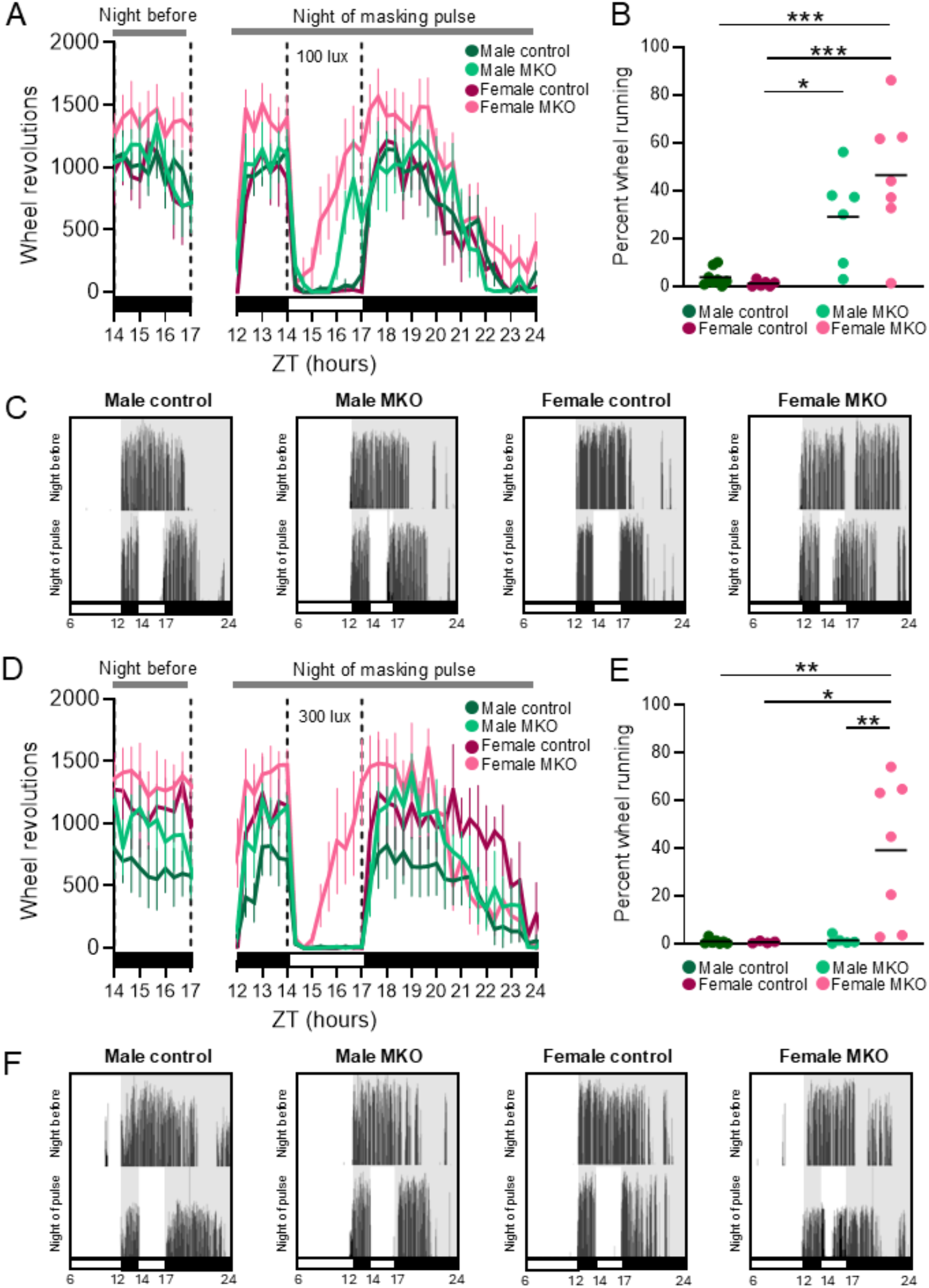
Lack of melanopsin results in a sex-specific deficit in negative masking. (A) Total wheel revolutions in 20-minute bins over the course of the night of a 3-hour, 100-lux negative masking pulse (denoted by white bar) from ZT14-17, as well as the same 3-hour period the night before. (B) Activity during the during the 100-lux light pulse as percent wheel revolutions compared to the same 3-hour period the night before. (Two-way ANOVA; Sex, F(1, 23) = 1.309, p = 0.2644; Genotype, F(1, 23) = 29.67, p < 0.0001, Šídák’s multiple comparisons test; Control Female vs MKO Male, p = 0.0486; Control Female vs MKO Female, p = 0.0004; Control Male vs MKO Female, p = 0.0003). (C) Example actograms of the night before and night of the 100-lux negative masking pulse. (D) Same as (A) but for a 300-lux negative masking pulse. (E) Same as (B) but for a 300-lux negative masking pulse. (Two-way ANOVA; Sex, F(1, 19) = 6.652, p = 0.0184; Genotype, F(1, 19) = 7.206, p = 0.0147, Šídák’s multiple comparisons test; Control Male vs MKO Female, p = 0.0028; Control Female vs MKO Female, p = 0.0108; MKO Male vs MKO Female, p = 0.0072). (F) Same as (C) but for a 300-lux negative masking pulse.

## Discussion

Here we identify previously unreported sex differences in retinal melanopsin expression and the role of melanopsin in photic phase delay and negative masking behavior. Interestingly, a loss of melanopsin *abolishes* a sex difference in phase shifting but *reveals* a sex difference in negative masking. For phase shifting, higher melanopsin expression in females drives larger light-induced phase delays than in males, while for negative masking melanopsin is necessary to achieve similar suppression of activity compared to males. Overall, the greater severity of circadian behavioral phenotypes in female MKO animals compared to males indicates that melanopsin phototransduction plays a larger role in shaping female light-driven circadian behaviors.

The SCN acts as the central circadian pacemaker in mammals, making it the primary focus of previous mechanistic exploration into the origin of sex differences in circadian behavior. Indeed, previous studies have reported sex differences in SCN neuronal morphology, neurochemistry, and function.^8^ Moreover, estrogen is known to modulate SCN neuron function^15,27^ and loss of the ESR1 estrogen receptor or ovariectomy both abolish previously reported sex differences in circadian behavior,^10,11,14^ indicating that hormonal modulation also plays a role in modulating these behaviors. Our results suggest that photoreception in ipRGCs is sex-specific and also plays a key role in shaping these same behaviors. Understanding how the sex difference in the expression of melanopsin that we report here affects SCN responses to light, and how distinct circuit and cellular-level sex differences combine to drive behavioral sex differences are important questions for future studies.

It is important to note that in addition to their indispensable role in circadian behaviors, ipRGCs also project widely to other brain nuclei involved in a multitude of processes like cognition and hormonal regulation,^29–31^ and have been shown to mediate light’s effects on a multitude of other behaviors and physiological functions, including sleep, mood, learning, memory, and anxiety.^32– 38^ Therefore differences in ipRGC function in males and females could result in broad downstream differences in health outside of circadian impacts, particularly for those exposed to aberrant lighting conditions like jet lag and shift work. Thus, understanding the mechanisms by which environmental light affects these health outcomes differently in males and females is crucial to developing effective preventions and treatments.

Overall, our results show the importance of directly comparing males and females in studies of light driven behaviors and retinal function. The novel sex difference in negative masking that we report here suggests that even for historically well-studied circadian behaviors, there are likely more circadian sex differences to be found, likely originating in the retina. Given that circadian disruption is linked to numerous negative health outcomes including cancer, obesity, cardiovascular disease, and reduced fertility, understanding these mechanisms has real potential to better align our depth of understanding of the male and female circadian systems to maximize potential positive health outcomes.^1–3,5,6,28^

### Limitations of the study

The role of melanopsin in driving behaviors is light intensity-dependent, as evidenced by our photic phase shifting results. Thus, one limitation of the current study is that we only tested 1-2 lighting intensities for each circadian behavior. Indeed, this may be the reason for which we did not see any sex differences in jet lag re-entrainment.

Another limitation of this study is that we did not track the estrous cycle of the female mice during behavioral testing in order to avoid introducing stress or disturbing their activity with daily vaginal lavages. While cycling estrogen certainly plays a role in shaping the behaviors that we tested, our results showing that melanopsin also plays a role are clear even in the without staging estrous.

## Resource Availability

### Lead Contact

Further information and requests for resources should be directed to the lead contact, Tiffany Schmidt

### Materials availability

This study did not generate new unique reagents.

### Data and code availability

Data can be obtained from the corresponding author T.M.S. upon reasonable request.

## Acknowledgments

This work has been supported by NIH grant F31 EY034387 to K.C.M., NSF grant DGE-1842165 to K.C.M., NIH grant 5R01GM143545-04 to J.A.E., and NIH grant 5R01EY030565-05 to T.M.S.

## Author contributions

K.C.M., J.A.E., and T.M.S. developed this study’s concept and design; K.C.M., M.L.A., J.D.B., and M.D. collected and analyzed data. K.C.M. and T.M.S. drafted the article and M.L.A, J.D.B., and J.A.E reviewed the article.

## EXPERIMENTAL MODEL AND SUBJECT DETAILS

All procedures were approved by the Animal Care and Use Committee at Northwestern University. We used male and female adult mice for all experiments. Mouse sex was determined by external anatomy. Control mice were Opn4^Cre/+ 39^ (RRID:IMSR_JAX:035925) and melanopsin null mice were Opn4^Cre/Cre^.

## METHOD DETAILS

### RNAscope

Eyes were enucleated and retinas dissected in PBS. Retinas were fixed in a 4% paraformaldehyde (PFA) solution in PBS for 24 hours at 4°C. Retinas were then washed in PBS followed by dehydration in methanol (MeOH) in PBS series (50%, 75%, and 100%) at room temperature for 5 minutes each. Retinas were stored in 100% methanol at -20°C overnight.

Retinas were re-hydrated in reverse MeOH in PBS steps and washed in PBS three times for 5 minutes each. Retinas were then incubated with RNAscope Protease Plus Reagent (Advanced Cell Diagnostics) for 30 minutes at 40°C, washed, and then incubated with probe for *Opn4* overnight. Retinas were washed and post-fixated for 10 minutes at room temperature in 4% PFA, washed, and then processed according to the RNAscope Multiplex Fluorescent v2 assay (Advanced Cell Diagnostics) instructions provided by the manufacturer. Tissue was mounted and sealed using ProLong Glass Antifade mountant (Thermo Fisher Scientific). Retinas were imaged on a Leica SP5 confocal microscope. For quantification, high magnification images (183.69 µm x 183.69 µm with a pixel size of 0.36 µm) with a z-stack size of 1 µm were taken.

ROIs around *Opn4* mRNA and control background were manually drawn and the mean of pixel intensity, per ROI, per stack, per probe was established using ImageJ. Images were processed, the mean background was subtracted, and label intensity data for each ROI on every whole mount retina was normalized against the maximum intensity value observed within that retina.

### Quantitative PCR

Mice were euthanized at one of four circadian time points and retinas were dissected in cold, nuclease-free PBS. Euthanasia and dissections for ZT 0 and ZT 18 were performed in the dark and those for ZT 6 and ZT 12 were performed in the light. Both retinas from each animal were pooled into one sample. Retinas were transferred to 1.5mL Eppendorf tubes, excess PBS was removed, and then 500μL of Trizol was added to each sample before homogenization and incubation at room temperature for 5 minutes. Next, 100μl chloroform was added to each sample before 3 minute incubation at room temperature and then centrifugation at 14,000 rpm for 15 minutes at 4°C. The resulting aqueous phase of each sample was transferred to a new tube. RNA was precipitated with the addition of 500μL of isopropanol and vortexed for 30 seconds. Samples were incubated at room temperature for 10 minutes with periodic vortexing. Samples were then centrifuged at 14,000 rpm for 15 minutes at 4°C, and the supernatant discarded. The remaining RNA pellet was washed by adding 75% ethanol in nuclease-free water and vortexing. Each sample was then centrifuged at 10,000 rpm for 5 min at 4°C. The supernatant was discarded and pellets left to air dry for 5 minutes. RNA pellets were then dissolved in 25μL of nuclease-free water. The RNA concentration of each sample was measured with a NanoDrop spectrophotometer (Thermo Fisher Scientific). cDNA was generated and then qRT-PCR was done using SYBR green and custom primers. Expression of melanopsin was normalized to the geometric mean of two housekeeping genes, *Gapdh* and *Hgprt*, in each sample.

### Wheel running

Mice were individually housed with ad libitum access to food and water in cages with running wheels and activity was recorded with ClockLab Data Collection software (Actimetrics, now Lafayette Life Sciences). Mice began in a 12:12 light dark (LD) cycle with 100-lux light during the light phase. Mice that ran inconsistently or were not well entrained were excluded.

For the first phase response experiment, mice were released into constant dark (DD). On the second day of DD, a 15-minute, 1000-lux pulse of light was given at CT14. The phase delay was calculated as the difference in hours between the onset of activity between the day after the pulse and the day of the pulse prior to the pulse. Onset of activity was manually marked on actograms by an experimenter blinded to genotype and sex.

For the second phase response experiment, mice were released into constant dark (DD) and on the third day of DD, a 1-hour, 150-lux pulse of light was given at CT16. Regression lines were fit for the onset of activity during the first 3 days of DD as well as the subsequent 6 days after the light pulse. The differences between these two lines was calculated on the day immediately following the light pulse using Clocklab.

For the jet lag experiment, mice were exposed to a 6-hour phase advance. Onset during re-entrainment was manually marked on actograms in ClockLab Analysis (Actimetrics, now Lafayette Life Sciences) by an experimenter blinded to genotype and sex.

For the negative masking experiments, a 3-hour, 100-lux or 300-lux light pulse was given from ZT14 to ZT17. Activity was binned into 20-minute bins. Masking was quantified as the wheel revolutions measured during the masking pulse as a percentage of the wheel revolutions during the same time the night before.

### Statistical comparisons

All graphs and statistical analyses were performed using Graph Pad Prism (RRID: SCR_002798).

---

**Figure S1.**
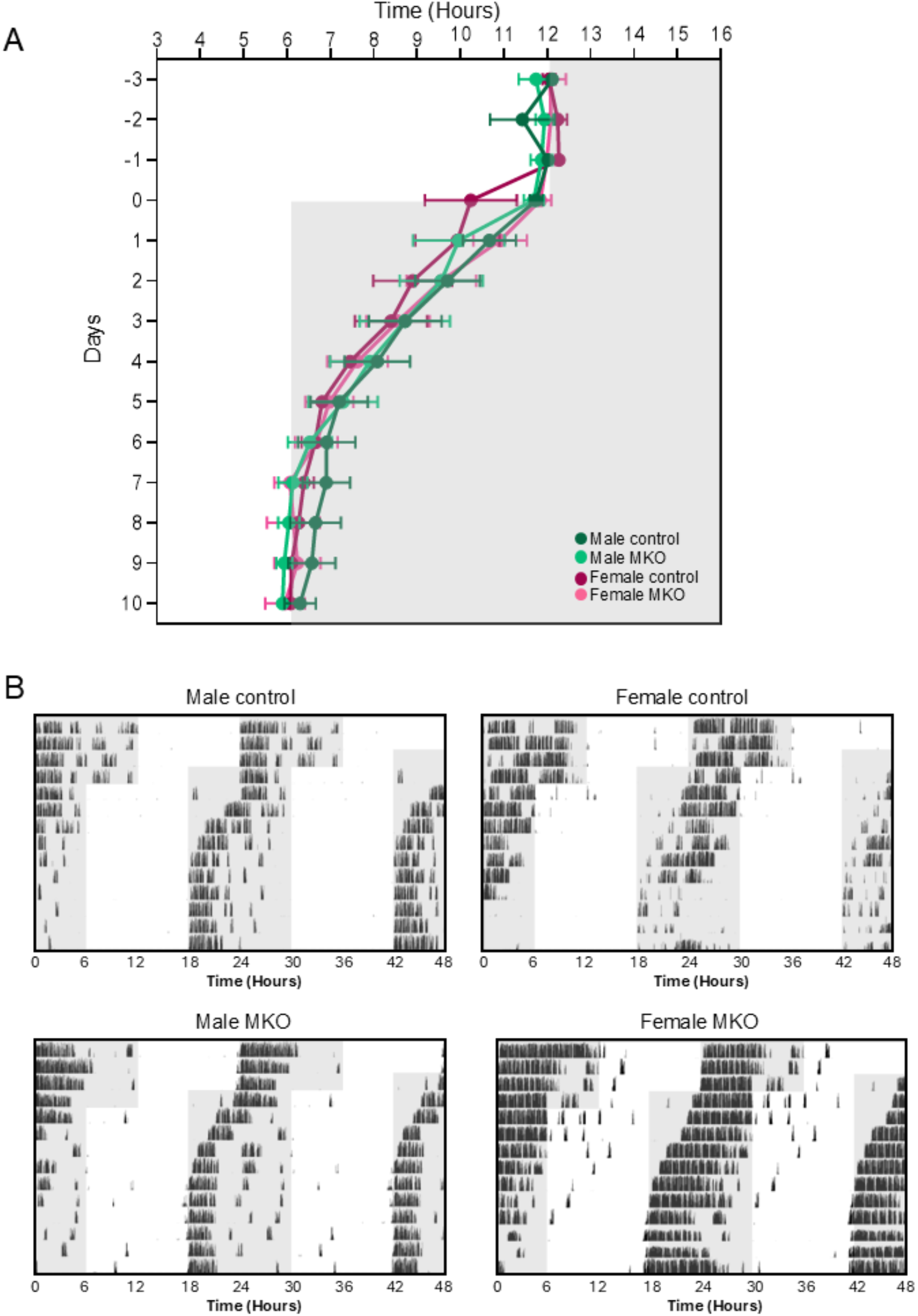
No melanopsin dependence or sex difference in jet lag advance. (A) Average onset of activity before and after a 6-hour phase advance on Day 0. Error bars indicate ± SEM. (B) Example actograms of activity during the 6-hour phase advance.

